# A neuronal social trait space for first impressions in the human amygdala and hippocampus

**DOI:** 10.1101/2021.01.30.428973

**Authors:** Runnan Cao, Chujun Lin, Xin Li, Alexander Todorov, Nicholas J Brandmeir, Shuo Wang

## Abstract

People instantaneously evaluate faces with significant agreement on evaluations of social traits. However, the neural basis for such rapid spontaneous face evaluation remains largely unknown. Here, we recorded from 490 neurons in the amygdala and hippocampus in 5 neurosurgical patients and show that amygdala and hippocampal neurons encode a social trait space. We further investigated the temporal evolution and modulation on the social trait representation, and we employed encoding and decoding models to reveal the critical social traits for the trait space. We also recorded from another 259 neurons and replicated our findings using different social traits. Lastly, the neuronal social trait space may have a behavioral consequence likely involved in the abnormal processing of social information in autism. Together, our results suggest that there exists a neuronal population code for a comprehensive social trait space in the human amygdala and hippocampus that underlie spontaneous first impressions.

## Introduction

Faces are among the most important visual stimuli we perceive and they often convey a wealth of information. When we see a person’s face, we can easily recognize their unique identity and general features such as sex and age. The gestalt of facial processing enables us to automatically evaluate faces on multiple trait dimensions (e.g., trustworthiness) (Willis & Todorov 2006), and these evaluations predict important social outcomes, ranging from electoral success to sentencing decisions (Todorov et al 2015). However, a central challenge in face research is to understand how the brain evaluates faces in general and forms rapid spontaneous impressions of faces on multiple trait dimensions.

It has been conclusively shown that neurons in the primate inferotemporal (IT) cortex encode a face space of low-level features, demonstrating a comprehensive neural code for physical variations in faces such as eye shape and skin tone (Chang & Tsao 2017, Freiwald et al 2009, Leopold et al 2006). On the other hand, the human amygdala and hippocampus play critical roles in social perception (Montagrin et al 2018, Rutishauser et al 2015) and encode various social trait judgments of faces (i.e., judgments of an individual’s temporally stable characteristics). For example, a lesion study has shown that the amygdala is necessary for judging facial trustworthiness (Adolphs et al 1998), which is further supported by functional neuroimaging studies (Todorov et al 2008). We previously utilized single-neuron recordings in the human amygdala to show that the amygdala parametrically encodes facial emotions (Wang et al 2017), which are known to shape various social trait judgments of faces such as personality traits (Said et al 2009). Prior studies have investigated one trait judgment at a time; however, humans use hundreds of different trait words to describe spontaneous trait judgments of faces (Lin et al 2019, Oosterhof & Todorov 2008, Sutherland et al 2013) and automatically evaluate faces on multiple trait dimensions simultaneously. Whether the amygdala and hippocampus encode a comprehensive space for social trait judgments of faces has not yet been determined.

In this study, we hypothesize there exists a neuronal social trait space in the human amygdala and hippocampus that underlies spontaneous first impressions of faces. Primate research on face processing supports such a possibility: neurons from the macaque temporal lobe encode a multi-dimensional face feature space (Chang & Tsao 2017, Freiwald et al 2009, Leopold et al 2006) as well as a multitude of social information (for a review see (Freiwald et al 2016)), providing plausible neural mechanisms supporting different dimensions of complex social evaluations. Furthermore, our recent neuroimaging data suggests that the human amygdala encodes physical variations in faces (e.g., shape and skin tone) that underlie representations of various social traits (Cao et al 2020a). A recent psychological study using the largest number of representatively sampled social traits to date has characterized a comprehensive space for social trait judgments of faces—a four-dimensional space with dimensions interpreted as warmth, competence, femininity, and youth (Lin et al 2019). Based on this comprehensive social trait space, the present study investigated whether there exists a population code (i.e., neuronal population activity collectively contributes to the judgments) for evaluating multimodal social traits in the human amygdala and hippocampus, which will provide the neural basis for first impressions of faces. We also provide a direct replication of our results using an additional dataset and another well-established social trait space. We lastly explore the behavioral consequence of the neuronal social trait space for those with autism.

## Results

### Constructing a comprehensive social trait space

Neurosurgical patients undergoing single-neuron recordings viewed 500 natural face images of 50 celebrities (**Figure 1A, B**; 10 images per celebrity) while performing a simple one-back task (**Figure 1A**; accuracy = 75.7±5.28% [mean±SD across sessions]). Additionally, we acquired consensus social trait ratings for the same face stimuli on 8 traits from a large population of participants recruited via an online platform (see **Methods**; 415.75±11.42 [mean±SD] raters per trait; **Figure S1A**). The eight traits (*warm*, *critical*, *competent*, *practical*, *feminine*, *strong*, *youthful*, and *charismatic*) were selected to represent the four comprehensive psychological dimensions of social trait judgments of faces (warmth, competence, femininity, and youth; two traits per dimension; see **Methods**) (Lin et al 2019). The inter-rater consistency of these ratings (**Figure S1B, C**) was comparable to the established study (Lin et al 2019) (see also **Figure S1D** for correlations between ratings from different modules [each module contained one face image per identity] and **Figure S1E** for correlations between social traits). We used the average ratings across participants per face on the eight traits to construct a “social trait space” (**Figure 1B**). We verified that this social trait space reproduced the four comprehensive dimensions of facial social trait judgments found in the prior study (Lin et al 2019) (**Table S1**). We found that this social trait space demonstrated an organized structure after projecting it onto a two-dimensional space for visualization using t-distributed stochastic neighbor embedding (t-SNE): different images of the same person were clustered, and the two t-SNE dimensions showed the change in two of the four comprehensive psychological dimensions (warmth and femininity) as expected. The trait judgment was highly consistent for different images of the same person (**Figure 1B** and **Figure S2A**). It is worth noting that we also collected social trait ratings from a subset of neurosurgical patients and we found that ratings from patients were generally consistent with the consensus ratings from the online sample (**Figure S1F, G**). Therefore, we used the consensus ratings for further analysis.

**Figure 1.**
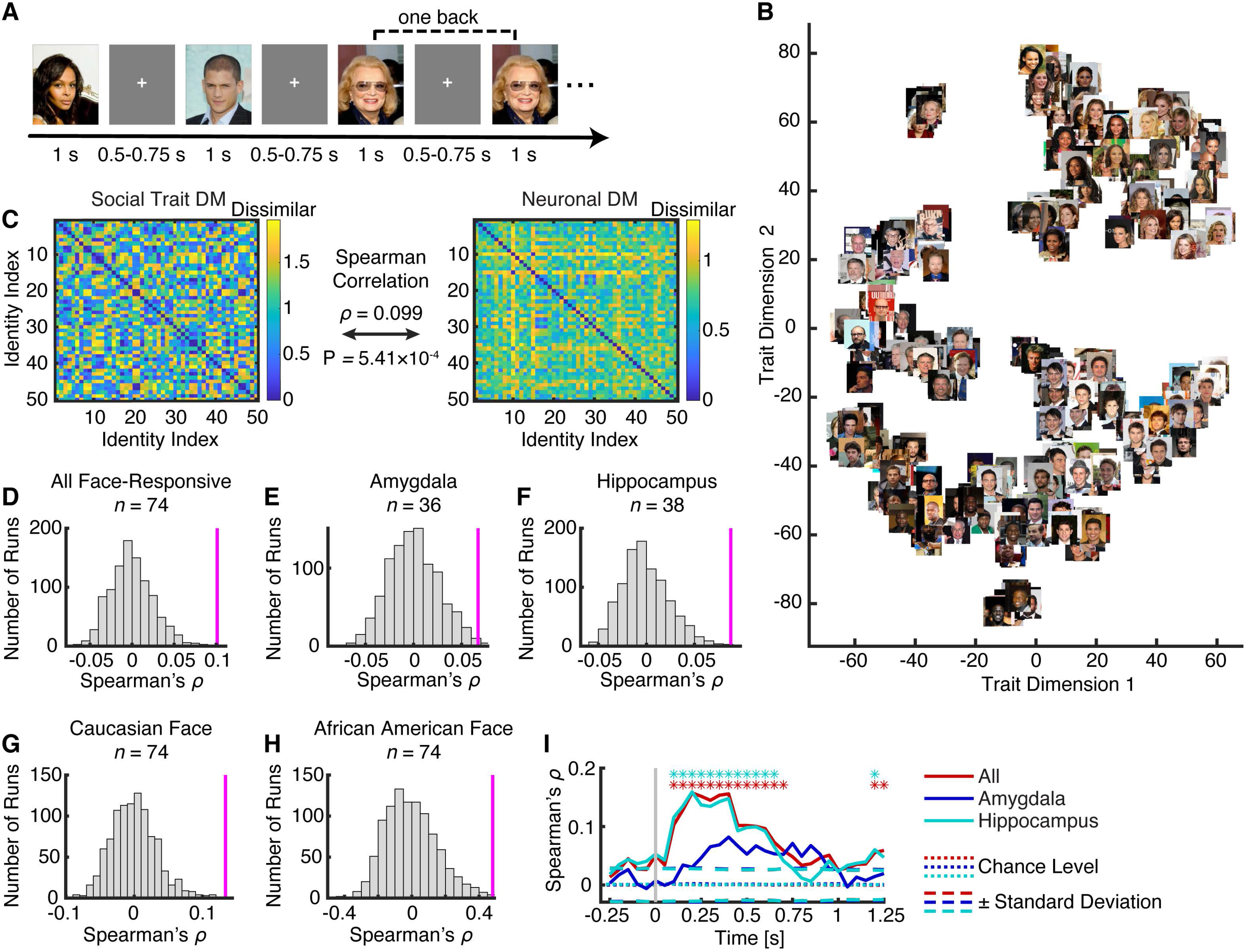
A neuronal social trait space. **(A)** Task. We employed a one-back task, in which patients responded whenever an identical face stimulus was repeated. Each face was presented for 1s, followed by a jittered inter-stimulus-interval (ISI) of 0.5 to 0.75 s. **(B)** Distribution of face images in the social trait space based on their consensus social trait ratings after dimension reduction using t-distributed stochastic neighbor embedding (t-SNE). **(C)**Correlation between dissimilarity matrices (DMs). The social trait DM (left matrix) was correlated with the neural response DM (right matrix). Color coding shows dissimilarity values. **(D-H)** Observed vs. permuted correlation coefficient between DMs. The correspondence between DMs was assessed using permutation tests with 1000 runs. The magenta line indicates the observed correlation coefficient between DMs. The null distribution of correlation coefficients (shown in gray histogram) was calculated by permutation tests of shuffling the face identities. **(D)** All face-responsive neurons (*n* = 74). **(E)** Amygdala face-responsive neurons (*n* = 36). **(F)** Hippocampal face-responsive neurons (*n* = 38). **(G)** Social trait space constructed by Caucasian faces only (*n* = 74). **(H)** Social trait space constructed by African American faces only (*n* = 74). **(I)** Temporal dynamics of correlation between DMs. Bin size is 500 ms and step size is 50 ms. The first bin is from −500 ms to 0 ms (bin center: −250 ms) relative to stimulus onset, and the last bin is from 1000 ms to 1500 ms (bin center: 1250 ms) after stimulus onset. Dotted horizontal lines indicate the chance level and dashed horizontal lines indicate the ±Standard Deviation (SD) of the null distribution. The top asterisks illustrate the time points with a significant correlation between DMs (permutation test against null distribution, P < 0.05, corrected by false discovery rate [FDR] Q < 0.05). See also **Figure S1** and **S2**.

### The neuronal population in the amygdala and hippocampus encode the social trait space

We recorded from 490 neurons in the amygdala and hippocampus of 5 neurosurgical patients (16 sessions in total; overall firing rate greater than 0.15 Hz), which included 242 neurons from the amygdala, 186 neurons form the anterior hippocampus, and 62 neurons from the posterior hippocampus. We aligned neuronal responses at stimulus onset and used the mean normalized firing rate in a time window from 250 ms to 1000 ms after stimulus onset for subsequent analyses.

To investigate whether the neuronal population encoded the comprehensive social trait space, we calculated dissimilarity matrices (DMs) between face identities for social traits (**Figure 1C** left; using ratings for 8 social traits) and neural responses (**Figure 1C** right; using the mean normalized firing rate of neurons), and we assessed the correspondence between the social trait DM and the neural response DM using representational similarity analysis (RSA) (Kriegeskorte et al 2008). We found that the DM from face-responsive neurons (i.e., neurons that had a significant change in firing rate after stimulus onset compared to baseline; see **Methods**; *n* = 74) was significantly correlated with the social trait DM (**Figure 1D**; permutation P < 0.001), and this was the case for both amygdala neurons (**Figure 1E**; *n* = 36, permutation P = 0.011) and hippocampal neurons (**Figure 1F**; *n* = 38, permutation P = 0.004). Furthermore, we observed similar results using the entire neuronal population (*n* = 490, permutation P = 0.003; amygdala neurons: *n* = 242, permutation P = 0.12; hippocampal neurons: *n* = 248, permutation P < 0.001). Although identity neurons (i.e., neurons that selectively encoded certain identities) (Cao et al 2020b) might enhance the correlation between face identities, we derived similar results when we excluded identity neurons (*n* = 57, permutation P = 0.004). Lastly, we derived similar results when we constructed DMs using face images instead of face identities (**Figure S2**; all permutation Ps < 0.008).

We further investigated the impact of race on social trait perceptions. We found that the amygdala and hippocampal neurons encoded the social trait space constructed with Caucasian faces only (**Figure 1G**; permutation P < 0.001) or African American faces only (**Figure 1H**; permutation P = 0.003), suggesting that encoding of social traits in the amygdala and hippocampus was independent of racial differences. In addition, we investigated the time course of the correspondence between the social trait DM and the neural response DM (**Figure 1I**). We found that encoding of the social trait space peaked at between 200 ms to 400 ms after stimulus onset. The response from hippocampal neurons peaked earlier (at ~150 ms; in comparison to ~400 ms for amygdala neurons) and was greater than that from amygdala neurons, but amygdala neurons had a more sustained response than hippocampal neurons (**Figure 1I**).

### Encoding and decoding models corroborate the RSA results and further reveal the critical social traits for the trait space

We next constructed encoding and decoding models to investigate the relationship between neural response and each individual social trait. Using an encoding model (see **Methods**), we identified subsets of neurons that significantly tracked social trait judgments (Pearson correlation between the mean normalized firing rate and the mean *z*-scored social trait ratings across 50 identities; see **Figure 2A** for an example of the trait *strong* and **Figure S3A** for a summary of the number of significant neurons for each trait). At the population level, we found that face-responsive neurons significantly encoded the judgments on the social traits associated with three comprehensive dimensions (warmth, competence, and femininity dimensions: *warm*, *critical*, *competent*, *practical*, *feminine*, and *strong*; **Figure 2B**; two-tailed one-sample *t*-test of correlation coefficient *r* against 0); and we observed similar results in all neurons (**Figure S3B**), all amygdala neurons (**Figure S3C**), and all hippocampal neurons (**Figure S3D**). Encoding of the social traits associated with the fourth comprehensive dimension (youth) was uniquely observed for the neural population in the amygdala (for *youthful* across face identities; **Figure S3C**). The encoding models using face images instead of face identities (Pearson correlation between the normalized firing rate and the *z*-scored social trait ratings across 500 face images) showed that the firing rate of all neurons significantly correlated with all four dimensions of social trait judgments (**Figure S3F**), three of the four dimensions in face-responsive neurons (**Figure S3E**) and all amygdala neurons (**Figure S3G**), and two of the four dimensions in all hippocampal neurons (**Figure S3H**). Furthermore, we found that the neural population encoded the first four principal components of the 8 social traits, which replicated the four comprehensive dimensions of social trait judgments of faces (Lin et al 2019). Lastly, we found similar results using the absolute value for Pearson’s *r* for each neuron (statistical significance was assessed using a permutation test).

**Figure 2.**
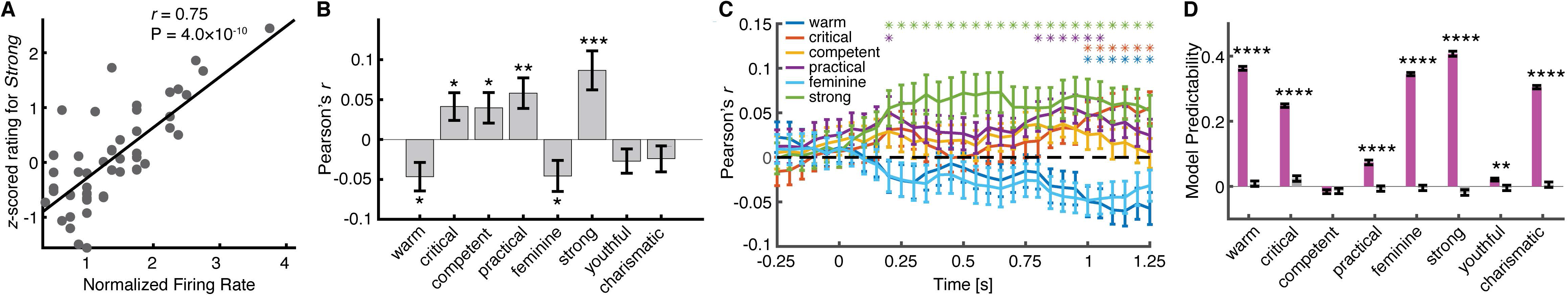
Encoding and decoding models. **(A)** An example neuron that showed a significant correlation between the mean normalized firing rate and the mean *z*-scored rating of the social trait *strong* (Pearson correlation: *r* = 0.75, P = 4.0×10^−10^). Each dot represents a face identity. **(B)** Encoding of each social trait. The bars show the average correlation coefficient across all face-responsive neurons for each social trait. Error bars denote ±SEM across neurons. Asterisks indicate a significant difference from 0 (two-tailed paired *t*-test). *: P < 0.05, **: P < 0.01 and ***: P < 0.001. **(C)**Encoding of different social traits over time. Error bars denote ±SEM across neurons. Asterisks shown on the top indicate a significant difference from 0 (two-tailed paired *t*-test, P < 0.05, corrected by false discovery rate [FDR] Q < 0.05). **(D)** Decoding of each social trait using a linear decoding model on face identities. Model predictability was assessed using the Pearson correlation between the predicted and actual trait ratings in the test dataset. The magenta bars show the observed response and the gray bars show the permuted response. Error bars denote ±SEM across permutation runs. Asterisks indicate a significant decoding performance (two-tailed two-sample *t*-test between observed vs. permuted). **: P < 0.01 and ****: P < 0.0001. See also **Figure S3** and **S4**.

Notably, we explored whether different social traits were encoded with a similar latency. To answer this question, we investigated the temporal dynamics of encoding models using a moving window. We found that the social trait associated with the femininity dimension regarding gender (*strong*) was encoded earlier after stimulus onset than the social traits associated with the warmth and competence dimensions (*warm*, *critical*, and *practical*) which describe the more abstract personality characteristics of an individual (**Figure 2C**) as opposed to physical characteristics.

Therefore, our results indicate that different social trait dimensions may be processed at different stages in the brain with physical characteristics being processed earlier than more complex personality traits. Furthermore, although amygdala and hippocampal neurons showed similar encoding for most of the social traits (**Figure S3C, D, G, H**; see **Figure S4** for temporal dynamics), we found that only amygdala neurons encoded the social trait *youthful*.

Using a decoding model (see **Methods**), we found that the neural population could predict the social trait judgments associated with all four dimensions (including the traits *warm*, *critical*, *practical*, *feminine*, *strong*, *youthful*, and *charismatic*) across face identities (**Figure 2D**) or across face images. Furthermore, we found similar results using partial least squares (PLS) regression (**Figure S3I**) and regression with principal component analysis (PCA) of neural responses (**Figure S3J**). Together, the encoding and decoding models corroborated our finding that neurons from the amygdala and hippocampus collectively encode a comprehensive social trait space.

### Social trait representation is universal for different face stimuli and social trait spaces

We conducted an additional experiment to (1) rule out the possibility that participants’ knowledge of some of the celebrities in our stimuli may influence neural representations of social traits, (2) investigate whether encoding of the social trait space can be generalized to different face stimuli and a social trait space constructed using a different set of social traits, and (3) explore whether encoding of the social trait space is independent of the evaluative context.

We recorded from a separate population of 259 neurons (12 sessions from 4 patients; firing rate > 0.15 Hz) while patients performed a trustworthiness judgement task (6 sessions; **Figure 3A**) or a dominance judgment task (6 sessions) using the FaceGen model faces (Oosterhof & Todorov 2008), which contained only feature information but no real identity information (**Figure 3A, B**). We used nine social traits (*attractiveness*, *competence*, *trustworthiness*, *dominance*, *mean*, *frightening*, *extroversion*, *threatening*, and *likability*) to construct a social trait space (**Figure 3B**). Similarly, we calculated DMs between faces for social traits (**Figure 3C**; using *z*-scored ratings for the 9 social traits) and neural responses (**Figure 3D-F**; using the mean normalized firing rate of neurons), and we assessed the correspondence between the social trait DM and the neural response DM using RSA. We found that the neural response DM was significantly correlated with the social trait DM (**Figure 3D, G**; permutation P < 0.001), suggesting encoding of the social trait space was independent of face familiarity, specific face stimuli (natural photos of real people vs. computer-generated model faces), and specific social traits to construct the space. Importantly, neurons encoded the social trait space separately in both the trustworthiness judgment task (**Figure 3E, H**; permutation P < 0.001) and the dominance judgment task (**Figure 3F, I**; permutation P = 0.010), suggesting that encoding of the social trait space was independent of the evaluative context. Together, this additional experiment not only confirmed our finding that neurons in the human amygdala and hippocampus encode a social trait space but also suggest that the social trait representation can be universal for different face stimuli and social trait spaces.

**Figure 3.**
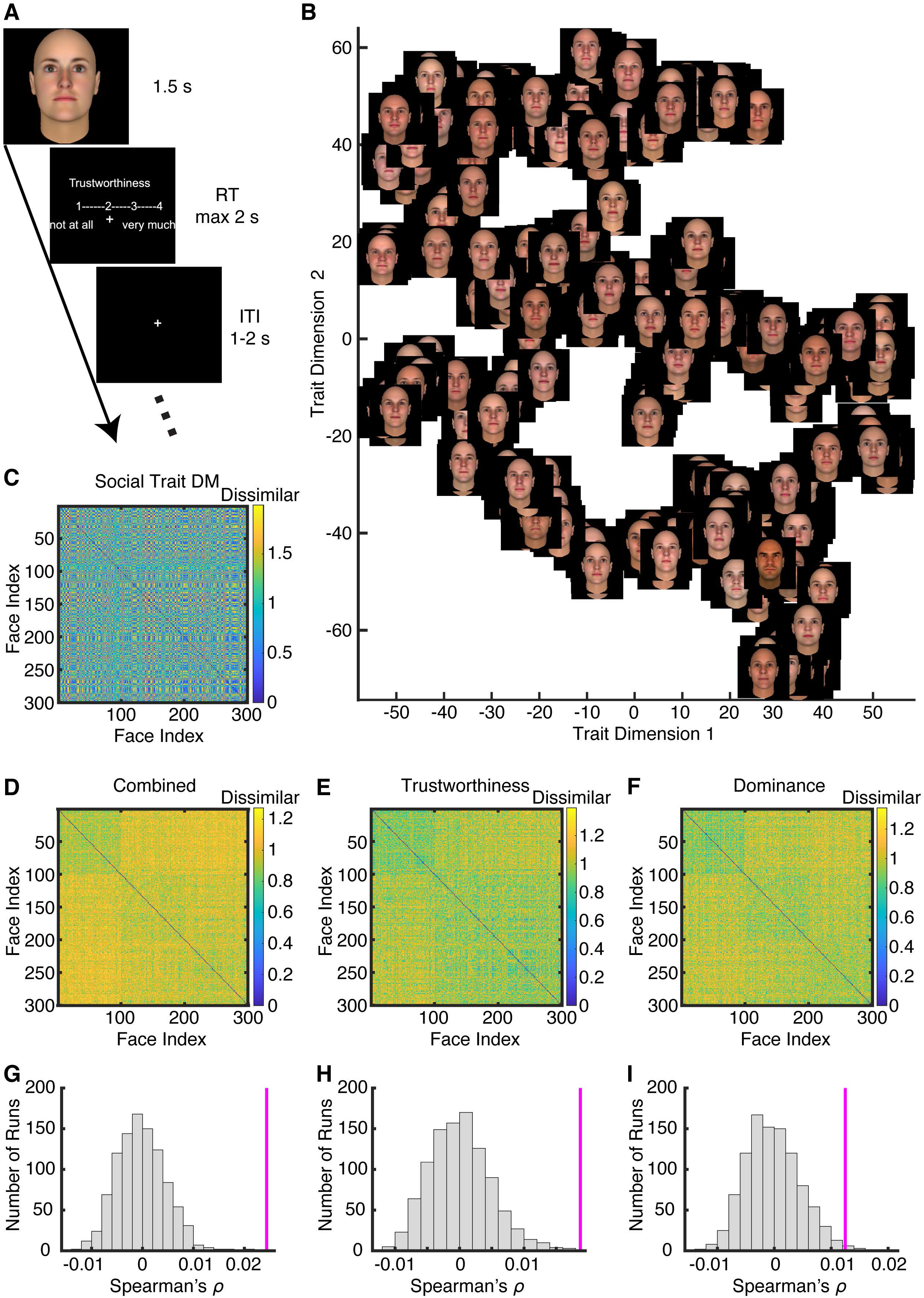
Replication of the neuronal social trait space using FaceGen stimuli and a different set of social traits. **(A)** Task. Each face was presented for 1.5 s, followed by participants’ judgment of trustworthiness / dominance within 2 s. The inter-trial-interval (ITI) was jittered between 1 to 2 s. **(B)** Distribution of face images in the social trait space based on their consensus social trait ratings after dimension reduction using t-distributed stochastic neighbor embedding (t-SNE). **(C)** The social trait dissimilarity matrix (DM). **(D-F)** The neural response DMs. Color coding shows dissimilarity values. **(G-I)** Observed vs. permuted correlation coefficient between DMs. The magenta line indicates the observed correlation coefficient between DMs. The null distribution of correlation coefficients (shown in gray histogram) was calculated by permutation tests of shuffling the faces (1000 runs). **(D, G)** Combined trustworthiness and dominance judgment tasks. **(E, H)** Trustworthiness judgment task. **(F, I)** Dominance judgment task.

### People with autism spectrum disorder (ASD) show an altered social trait representation

People with ASD demonstrate abnormal processing of social information from faces (Adolphs et al 2001). A specific neural structure hypothesized to underlie deficits in face processing in ASD is the amygdala, a brain structure that has long been implicated in autism (Baron-Cohen et al 2000, Schumann & Amaral 2006). Therefore, in the present study we also explored whether people with ASD have a different social trait representation compared to controls and whether the social trait representation in ASD can be explained by neural response from the amygdala and hippocampus.

To address these questions, we acquired ratings of the CelebA stimuli from a sample of online participants with ASD (self-identified). We first confirmed that online participants with ASD demonstrated significantly higher scores compared to controls on standardized tests that evaluate ASD characteristics including the Autism Spectrum Quotient (AQ; **Figure 4A**; ASD: 27.76±8.09 [mean±SD], controls: 20.28±6.82; two-tailed two-sample *t*-test: *t*(427) = 8.94, P < 10^−16^) and Social Responsiveness Scale-2 Adult Self Report (SRS-A-SR; **Figure 4B**; ASD: 91.73±29.66, controls: 65.17±25.19; *t*(427) = 8.61, P = 1.11×10^−16^). In comparison, neurosurgical patients had scores comparable to controls for both AQ (**Figure 4A**; 18.25±8.46; *t*(339) = 0.59, P = 0.56) and SRS-A-SR (**Figure 4B**; 31.5±20.51; *t*(337) = 1.88, P = 0.06). We also confirmed that online participants had scores similar to well-characterized in-lab participants from our prior ASD study (Wang & Adolphs 2017a) for both AQ (ASD: *t*(108) = 0.73, P = 0.47; controls: *t*(349) = 1.42, P = 0.16) and SRS-A-SR (ASD: *t*(107) = 0.94, P = 0.35; controls: *t*(343) = 1.57, P = 0.16).

**Figure 4.**
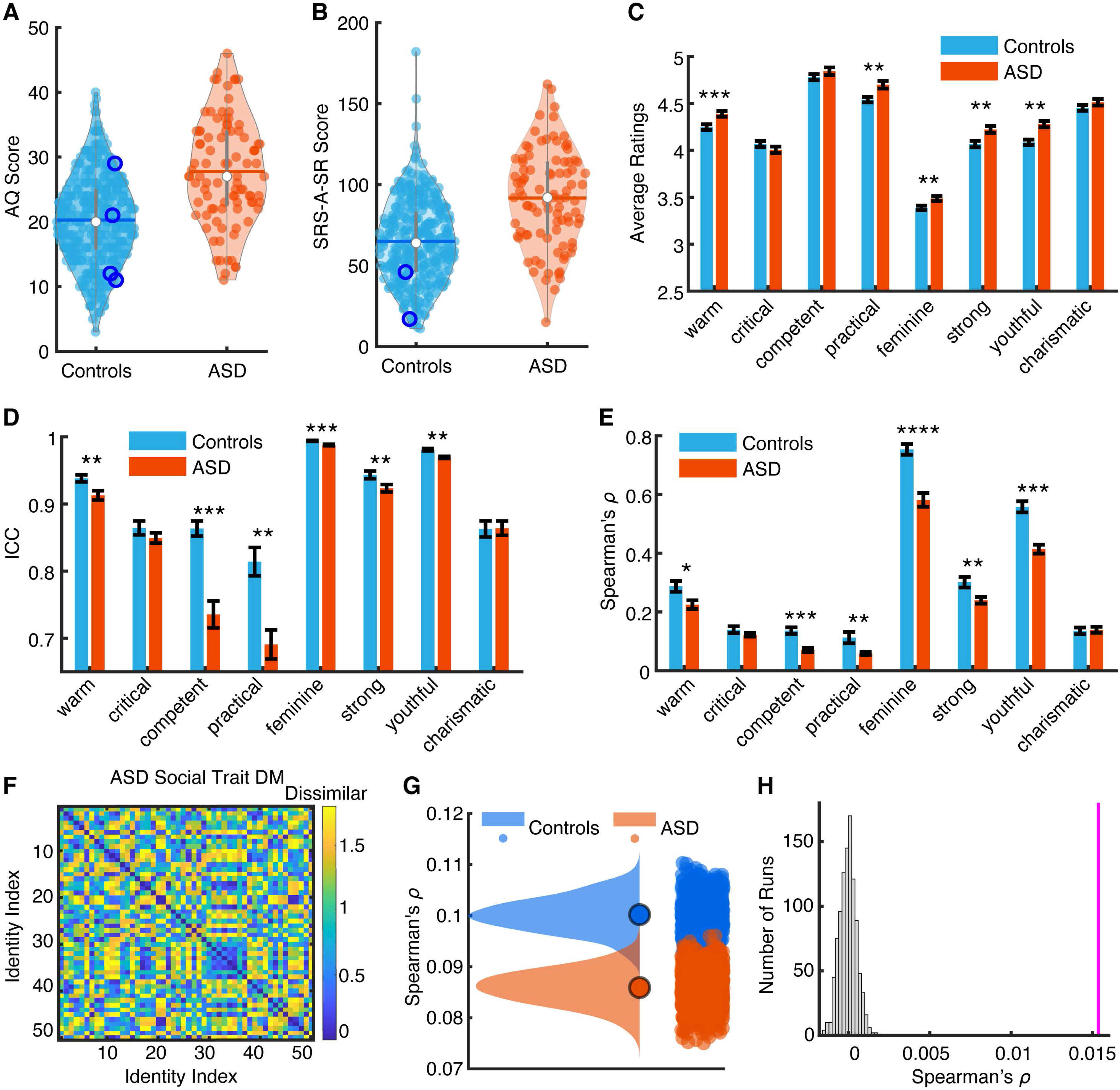
People with ASD demonstrate atypical social trait representations. **(A)** Autism Spectrum Quotient (AQ). **(B)** Social Responsiveness Scale-2 Adult Self Report (SRS-A-SR). Violin plots present the median value as the white circle and the interquartile range as the gray vertical bars. Blue circles show the scores from neurosurgical patients, which are comparable to those from controls. **(C)** Social judgment rating for each trait. **(D, E)** Inter-rater consistency. Inter-rater consistency of each trait was estimated using **(D)** the intraclass correlation coefficient (ICC) (McGraw & Wong 1996) and **(E)** the Spearman’s correlation coefficient (*ρ*). Inter-rater consistency was first calculated between raters and averaged within each module, and then averaged across modules. Error bars denote ±SEM across rating modules. Asterisks indicate a significant difference between participants with ASD and controls using two-tailed paired *t*-test. *: P < 0.05, **: P < 0.01, ***: P < 0.001, and ****: P < 0.0001. **(F)** The social trait dissimilarity matrix (DM) from participants with ASD. **(G)** Bootstrap distribution of DM correspondence for each participant group. Blue: controls. Red: ASD. The dots show the mean value of each distribution. Participants with ASD showed a weaker correspondence with the neural response DM compared to controls. **(H)** Observed vs. permuted difference in DM correspondence between participant groups. The magenta line indicates the observed difference in DM correspondence between participant groups. The null distribution of difference in DM correspondence (shown in gray histogram) was calculated by permutation tests of shuffling the participant labels (1000 runs).

We found that the social trait ratings differed in all four comprehensive dimensions (including the traits *warm*, *practical*, *feminine*, *strong*, and *youthful*) between participants with ASD and controls (**Figure 4C**). Furthermore, participants with ASD demonstrated a significantly lower inter-rater consistency compared to controls in most of the traits (**Figure 4D, E**). This suggests that participants with ASD were more variable in their ratings, consistent with the heterogeneity in their symptoms and behavior (Happe et al 2006). Notably, although the social trait DM for participants with ASD (**Figure 4F**) was similar to controls, it was less correlated with the neural response DM from the neurosurgical patients (derived with face-responsive neurons; *ρ* = 0.084 for ASD and *ρ* = 0.10 for controls; similar results were derived with all neurons). We used a bootstrapping approach to estimate the distribution of DM correspondence for each participant group (see **Methods**) and we found that the two distributions were largely separated (**Figure 4G**; the mean of the ASD distribution was significantly outside the control distribution [P < 0.001] and the mean of the control distribution was also significantly outside the ASD distribution [P < 0.001]). We also used a permutation test (see **Methods**) and statistically confirmed that the difference in DM correspondence between participant groups was above chance (**Figure 4H**; P < 0.001). Together, our results suggest that the neuronal social trait space in the amygdala and hippocampus may have a behavioral consequence for social judgment and may account for abnormal social trait judgments of faces in ASD.

## Discussion

Our present results represent the first step towards constructing a social trait space for face processing at the single-neuron level in the human brain. We not only showed that single neurons encoded individual social traits when judging photos of famous people, but also demonstrated that the neuronal population in the amygdala and hippocampus encoded a comprehensive social trait space. In addition, we had a direct replication of our results using unfamiliar faces and different social traits, and our results further suggested that the neuronal social trait space in the amygdala and hippocampus may relate to abnormal face processing in autism based on behavioral data on social traits of the same stimuli.

Past behavioral research has provided candidate dimensions for describing trait judgments of faces (Lin et al 2019, Oosterhof & Todorov 2008, Sutherland et al 2013); however, the biological bases of those psychological dimensions remain unknown. Here we showed that these dimensions were encoded by the neural population in the amygdala and hippocampus. We further showed that the neural correlates for different social trait dimensions varied in temporal dynamics (i.e., the femininity dimension [a physical characteristic] was encoded faster than the more abstract dimensions of warmth and competence [personality traits]), suggesting that different categories of social trait information may arrive at the amygdala and hippocampus at different latencies. This result is consistent with the notion that the amygdala connects with other parts of the brain through multiple routes (Pessoa & Adolphs 2010). In addition, we found that the dimensions of warmth, competence, and femininity were encoded across both face images and face identities, whereas the dimension of youth was only encoded across face identities regarding *youthful* and across face images regarding *charismatic*, likely because different face images from the same identity were more heterogenous along the youth dimension. Similar neural pattern analyses have been used to study race bias of faces using functional neuroimaging data (Stolier & Freeman 2016) and face representation using intracranial electroencephalogram data (Grossman et al 2019) in humans.

It is worth noting that our one-back task did not require patients to make any explicit face judgment (they simply indicated when a face was repeated); therefore, our analyses were relating neural responses of implicit face impressions provided by patients to the consensus ratings of explicit face impressions provided by an independent sample of over 400 participants from the general population. Using an additional experiment with computer-generated, unfamiliar faces, we further illustrated that encoding of the social trait space was independent of face familiarity, the knowledge of the face identity, as well as specific faces and traits being evaluated. Therefore, the neural coding in the amygdala and hippocampus can be a general mechanism for face evaluation and first impressions. Furthermore, it has been argued that the representation of social trait space is subject to top-down modulation (Stolier et al 2018), and our previous findings also suggested a flexible representation of social traits (e.g., trustworthiness and dominance) in the human amygdala (Cao et al 2020a). However, in our additional experiment (**Figure 3**), we showed that encoding of the social trait space was universal under different evaluative contexts. Also, in our main experiment (which likely involved both bottom-up and top-down processes) we found that the representation of social traits was similar for different races. Therefore, future research will be needed to investigate how the two distinct processes (bottom-up and top-down) may influence the neuronal representation of social traits.

The ability to look at someone’s face and make judgments about basic aspects of that person, such as trustworthiness, is a fundamental skill for most people. Unfortunately, people with ASD have difficulty in processing social information from faces (Adolphs et al 2001). Such deficits in ASD may be attributed to the amygdala, a hypothesis that is supported by substantial literature showing structural abnormalities (Amaral et al 2008) and atypical brain activation (Philip et al 2012) in ASD. Furthermore, patients with amygdala damage also fail to fixate on the eyes when viewing faces similar to those with ASD (Adolphs et al 2005), and single-neuron recordings in the human amygdala show weaker responses in people with ASD when viewing the eyes (Rutishauser et al 2013). In this study, we further showed that the neuronal social trait space encoded by amygdala and hippocampal neurons may account for aspects of abnormal face processing in ASD since behavioral ratings of social traits using the same stimuli by adults with ASD were less correlated with the neuronal data from neurosurgical patients than was the behavioral ratings of social traits from controls.

In conclusion, we for the first time revealed a neural social trait space at the single-cell level in humans. Encoding a comprehensive social trait space provides the neural basis for rapid spontaneous impressions of faces on multiple trait dimensions. Our present results are in line with the notion that face representations are encoded over a broad and distributed population of neurons (Hinton 1984), which has been conclusively demonstrated in the non-human primate inferotemporal cortex (IT) (Chang & Tsao 2017). Our results further shed light on how face processing evolves along the visual processing stream where the brain transforms from encoding low-level facial features in the higher visual cortex to complex social traits in the amygdala and hippocampus. Our results also support the idea that the amygdala and hippocampus are highly involved in social perception and evaluation (Adolphs 2010, Montagrin et al 2018), which in turn supports their roles in coding socially relevant and salient stimuli (Wang & Adolphs 2017b).

## Acknowledgements

We thank all patients for their participation, staff from WVU Ruby Memorial Hospital for support with patient testing, and Paula Webster for valuable comments. This research was supported by an NSF CAREER Award (1945230), Air Force Young Investigator Program Award (20RT0829), Dana Foundation Clinical Neuroscience Award, ORAU Ralph E. Powe Junior Faculty Enhancement Award, West Virginia University (WVU), and WVU PSCoR Program (to S.W.), and an NSF Grant (OAC-1839909) and the WV Higher Education Policy Commission Grant (HEPC.dsr.18.5) (to X.L.). The funders had no role in study design, data collection and analysis, decision to publish, or preparation of the manuscript.

## Author Contributions

R.C., C.L., X.L., and S.W. designed research. R.C. and S.W. performed experiments. N.J.B. performed surgery. R.C., C.L., and S.W. analyzed data. R.C, C.L., X.L., A.T., and S.W. wrote the paper. All authors discussed the results and contributed toward the manuscript.

## Methods

### Patients

A total of 16 single-neuron recording sessions were conducted with 5 patients (4 female) who had undergone surgery to have electrodes implanted to treat intractable epilepsy. All patients provided written informed consent using procedures approved by the Internal Review Board of West Virginia University (WVU).

### Stimuli

We used photos of celebrities from the CelebA dataset (Liu et al 2015). We selected 50 identities with 10 images for each identity, totaling 500 face images. The identities were selected to include both genders (33 male) and multiple races (40 identities were Caucasian, 9 identities were African American, and 1 identity was biracial). The faces were of different angles and gaze directions, with diverse backgrounds and lighting. The faces showed various facial expressions, with some having accessories such as sunglasses and hats. The same stimuli were used for all patients.

In addition, we used a validation dataset. We used the FaceGen Modeller program (http://facegen.com; version 3.1) to randomly generate 300 faces (see (Oosterhof & Todorov 2008) for detailed procedures). FaceGen constructs face space models using information extracted from 3D laser scans of real faces. To create the face space model, the shape of a face was represented by the vertex positions of a polygonal model of fixed mesh topology. With the vertex positions, a principal component analysis (PCA) was used to extract the components that accounted for most of the variance in face shape. Each principal component (PC) thus represented a different holistic non-localized set of changes in all vertex positions. The first 50 shape PCs were used to construct faces that had a symmetric shape. Similarly, because skin texture is also important for face perception, 50 texture PCs based on PCA of the RGB values of the faces were also used to represent faces. The resulting 300 faces were randomly generated from the 50 shape and 50 skin texture components with the constraint that all faces were set to be Caucasian. Notably, we have already acquired trait judgments of these faces from healthy control raters on 9 traits (Oosterhof & Todorov 2008): attractiveness, competence, trustworthiness, dominance, mean, frightening, extroversion, threatening, and likability. The trait judgements were measured on 9-point scales, ranging from 1 (not at all [trait]) to 9 (extremely [trait]). Therefore, these faces have benchmark ratings and we can readily perform correlational analysis with neural responses and psychometric behavioral data.

### Experimental procedure

We used a 1-back task for CelebA stimuli. In each trial, a single face was presented at the center of the screen for a fixed duration of 1 second, with uniformly jittered inter-stimulus-interval (ISI) of 0.5-0.75 seconds (**Figure 1A**). Each image subtended a visual angle of approximately 10º. A simple 1-back task required patients to press a button if the present face image was *identical* to the immediately previous image. Nine percent of the trials were one-back repetitions. Each face was shown once unless repeated in one-back trials; and the faces shown in one-back trials were randomly selected for each patient. We excluded responses from one-back trials to have an equal number of responses for each face. This task kept patients attending to the faces, but avoided potential biases from focusing on any particular facial feature (e.g., the color of their eyes or whether they are smiling) or social judgment (e.g., whether they seem happy). The order of faces was randomized for each patient.

For FaceGen stimuli, patients performed two face judgment tasks (**Figure 3A**). In each task, there was a judgment instruction, i.e., patients judged how trustworthy or how dominant a face was. We used a 1-4 scale: ‘1’: not trustworthy / dominant at all, ‘2’: somewhat trustworthy / dominant, ‘3’: trustworthy / dominant, and ‘4’: very trustworthy / dominant. Each image was presented for 1.5 s at the center of the screen.

Stimuli were presented using MATLAB with the Psychtoolbox 3 (Brainard 1997) (http://psychtoolbox.org) (screen resolution: 1600 × 1280).

### Online rating of social traits

For CelebA stimuli, we acquired social trait ratings of the faces from both patients and a large number of participants from the general population (age [M = 26.20 years, SD = 7.11], 180/500 females). Participants were asked to rate the faces on eight social traits using a 7-point Likert scale through an online rating task. The social traits included *warm, critical, competent, practical, feminine, strong, youthful,* and *charismatic*, representing the four core psychological dimensions of comprehensive trait judgments of faces (warmth, competence, femininity, and youth; 2 traits per dimension); and these social traits were well validated in a previous study (Lin et al 2019). The two traits per dimension were selected to capture the meaning of the dimension (e.g., *warm* for the warmth dimension) and they had the highest loadings for that dimension based on the prior data (Lin et al 2019) (e.g., *critical* had the highest loading on the warmth dimension). Furthermore, using more than one trait per dimension allowed us to perform principal component analysis (PCA) to validate our social trait space with one from the prior study (Lin et al 2019). Face stimuli used for collecting the consensus ratings were the same set of stimuli used for the neural recordings. Participants also indicated whether they could recognize the identity of the face (i.e., whether they were familiar with each face identity).

Patients completed the social trait rating task online after they were discharged from the hospital following surgery to treat intractable epilepsy. Three patients completed the rating task and provided ratings for 2 to 5 photos per identity per social trait. Participants from the general population completed the rating task using the Prolific online research platform. Participants who were fluent in English were included. We divided the experiment into 10 modules, with each module containing one face image randomly selected per face identity (totaling 50 face images per module), and we required participants to rate each face within a module on 8 social traits (e.g., competence; rated in blocks). We collected data from 50 participants per module. All face images in the stimulus set (500 in total) were rated on all 8 social traits, totaling 200,000 ratings across all participants and modules (500 face images x 8 traits x 50 participants).

We applied the following three exclusion criteria:

1. Trial-wise exclusion: we excluded trials with reaction times shorter than 100 ms or longer than 5000 ms.
2. Block/trait-wise exclusion: we excluded the entire block per participant if more than 30% of the trials were excluded from the block per (1) above, or if there were fewer than 3 different rating values in the block (this suggests that the participant may not have used the rating scale properly).
3. Participant-wise exclusion: we excluded a participant if more than 3 blocks were excluded from the participant per (2) above.

We subsequently applied a t-distributed stochastic neighbor embedding (t-SNE) method to convert the 8-dimensional social trait ratings into a 2-dimensional space for visualization (van der Maaten & Hinton 2008).

We repeated the same procedure to acquire ratings from participants with autism spectrum disorder (ASD). In addition, both participants with ASD and controls were asked to provide demographic information and complete the online Autism Spectrum Quotient (AQ) and Social Responsiveness Scale-2 Adult Self Report (SRS-A-SR) questionnaires.

### Inter-rater consistency

Inter-rater consistency of each trait was estimated using the intraclass correlation coefficient (ICC) (McGraw & Wong 1996) and the Spearman’s correlation coefficient (*ρ*). The ICC and Spearman’s *ρ* were computed between raters for each trait in each module and then averaged across modules per trait. The ICC was calculated using Matlab implementation written by Arash Salarian (2020) (https://www.mathworks.com/matlabcentral/fileexchange/22099-intraclass-correlation-coefficient-icc). The Spearman’s *ρ* was computed between each pair of raters and then averaged across all pairs of raters.

### Electrophysiology

We recorded from implanted depth electrodes in the amygdala and hippocampus from patients with pharmacologically intractable epilepsy. Target locations in the amygdala and hippocampus were verified using post-implantation CT. At each site, we recorded from eight 40 µm microwires inserted into a clinical electrode as described previously (Rutishauser et al 2006a, Rutishauser et al 2010). Efforts were always made to avoid passing the electrode through a sulcus, and its attendant sulcal blood vessels, and thus the location varied but was always well within the body of the targeted area. Microwires projected medially out at the end of the depth electrode and examination of the microwires after removal suggests a spread of about 20-30 degrees. The amygdala electrodes were likely sampling neurons in the mid-medial part of the amygdala and the most likely microwire location is the basomedial nucleus or possibly the deepest part of the basolateral nucleus. Bipolar wide-band recordings (0.1-9000 Hz), using one of the eight microwires as the reference, were sampled at 32 kHz and stored continuously for off-line analysis with a Neuralynx system. The raw signal was filtered with a zero-phase lag 300-3000 Hz bandpass filter and spikes were carefully sorted using a semi-automatic template matching algorithm as described previously (Rutishauser et al 2006b).

### Single-neuron response

Only units with an average firing rate of at least 0.15 Hz during the entire task were considered. Only single units were considered. Trials were aligned to stimulus onset. We used the mean firing rate in a time window 250 ms to 1250 ms after stimulus onset as the response to each face. Firing rate was then normalized by dividing the mean activity in the baseline (−250 ms to 0 ms relative to stimulus onset). Such normalization was applied in previous studies that analyzed the similarity between single-neuron responses to visual categories (Reber et al 2019).

Face-responsive neurons were identified by comparing the response to faces (i.e., the mean firing rate in a time window 250 ms to 1250 ms after stimulus onset) to baseline (i.e., −250 ms to 0 ms relative to stimulus onset) using a two-tailed paired *t*-test with P < 0.05.

### Representational similarity analysis (RSA)

Dissimilarity matrices (DMs) (Kriegeskorte et al 2008) are symmetrical matrices of dissimilarity between all pairs of face images or face identities. In a DM matrix, larger values represent larger dissimilarity of pairs, such that the smallest value possible is the similarity of a condition unto itself (dissimilarity of 0). We used the Pearson correlation to calculate DMs (ratings were *z*-scored and firing rates were normalized to the mean baseline of each neuron), and we used the Spearman correlation to calculate the correspondence between the DMs (Spearman correlation was used because it does not assume a linear relationship (Stolier & Freeman 2016)). We further used permutation tests with 1000 runs to assess the significance of the correspondence between the social trait DM and the neural response DM. Because the consistency between face images for the same face identity in both social trait ratings (**Figure S2A**) and neural responses (**Figure S2B**) could inflate the correspondence between the social trait DM and the neural response DM, we averaged the social trait ratings or neural responses across face images for each face identity and calculated the DM between face identities. We further used a moving window (bin size = 500 ms, step size = 50 ms) to measure temporal dynamics. The first bin started −500 ms relative to trial onset (bin center was thus 250 ms after trial onset), and we tested 31 consecutive bins (the last bin was thus from 1000 ms to1500 ms after trial onset).

We used a bootstrap with 1000 runs to estimate the distribution of DM correspondence for each participant group. In each run, 70% of the data were randomly selected from each participant group and we calculated the correspondence (Spearman’s ρ) between the social trait DM and the neural response DM for each participant group. We then created a distribution of DM correspondence for each participant group, and we compared the mean of the ASD distribution to the control distribution and vice versa to derive statistical significance.

We further used a permutation test with 1000 runs to statistically compare the DM correspondence between participants with ASD and controls. In each run, we shuffled the participant labels and calculated the difference in DM correspondence between participant groups. We then compared the observed difference in DM correspondence between participant groups with the permuted null distribution to derive statistical significance.

### Encoding and decoding models

We constructed encoding and decoding models to investigate the relationship between the neural response and social trait judgments. For the encoding model, we first calculated the Pearson correlation coefficient between each social trait (*z*-scored) and the response of each neuron (normalized by the baseline) across face images or face identities, and then we averaged the correlation coefficient *r* across neurons per social trait. We compared the mean correlation coefficient with 0 using a two-tailed one-sample *t*-test to determine if a social trait was significantly encoded by the neural population.

For decoding, we employed three models to linearly decode the social trait judgments from the neural response (note that the decoding was performed for each trait individually). First, we fitted a linear model for social trait ratings (*z*-scored) using neural responses (normalized by the baseline). We calculated the vector of neural weights *w* as: *w* = *R* **·** *v*, where *v* is a column vector (*n*×1) of trait ratings across the *n* faces or identities (*n* = 500 for faces and *n* = 50 for identities), and *R* is the neural matrix (each row is a neuron’s response and each column is a face or identity) that contains the neural response values for each face or identity. We further normalized *w* by || *w*||: *w* = *w* / ||*w*||. The resulting neural vector *w* thus showed the optimal direction that best captured the variation in trait ratings. Second, because there were more independent variables (i.e., neurons) than observations (i.e., trait ratings of identities), we employed a partial least squares (PLS) regression with 3 components to decode the social traits. Third, similarly, we first performed a principal component analysis (PCA) on the neural matrix and then used the first 3 principal components for a linear regression. To estimate statistical significance, for all three linear decoding models, we used a permutation test with 1000 runs to determine whether judgments of a social trait could be significantly decoded by the neural population response. In each run, we randomly selected 70% of the trait ratings as the training dataset to construct a decoding model (i.e., deriving neural model or regression coefficients). We then predicted the trait ratings for faces using this model in the test dataset, and computed the Pearson correlation between the predicted and actual trait ratings in the test dataset. As a reference, we randomly shuffled the trait ratings across faces to create a null distribution of Pearson correlations. A decoding model was considered *significant* if the mean correlation coefficient of the observed distribution was significantly greater than that of the null distribution (two-tailed two-sample *t*-test). This procedure has been demonstrated to be very effective in a recent study investigating neural population coding of facial features (Chang & Tsao 2017).

