## Supplementary Materials for "A neuronal social trait space for first impressions in the human amygdala and hippocampus"

**Table S1.** Factor loadings of the social trait space. The first column lists the social traits on which we collected consensus ratings, and the second row labels the four factors. We performed principal component analysis on the consensus social trait ratings across the 50 face identities (columns 2-5) and the 500 face images (columns 6-9). The first four principal components before rotation explained 66%, 19%, 8%, 5% of the variance in the data across face identities (total 98%; each of the remaining PCs explained less than 1% of the variance), and 59%, 18%, 11%, 7% of the variance in the data across face images (total 96%; each of the remaining PCs explained less than 2% of the variance). Varimax rotation was applied for interpretability of the factors, which replicated the four dimensions of comprehensive trait judgments of faces found in the prior research. The largest absolute factor loading across the four dimensions for each social trait space is highlighted in bold.

| | **Dimensions of the Social Trait Space (Across 50 Face Identities)** | | | | **Dimensions of the Social Trait Space (Across 500 Face Images)** | | | |
|---|---|---|---|---|---|---|---|---|
| **Social Traits** | Warmth | Competence | Femininity | Youth | Warmth | Competence | Femininity | Youth |
| *Critical* | **−0.94** | 0.23 | 0.07 | −0.11 | **−0.91** | 0.22 | 0.04 | −0.11 |
| *Warm* | **0.87** | 0.20 | 0.33 | 0.28 | **0.88** | 0.19 | 0.29 | 0.21 |
| *Charismatic* | **0.85** | 0.41 | 0.12 | 0.16 | **0.82** | 0.41 | 0.09 | 0.15 |
| *Competence* | 0.12 | **0.96** | 0.12 | −0.05 | 0.11 | **0.94** | 0.09 | −0.08 |
| *Practical* | 0.03 | **0.95** | −0.16 | −0.13 | 0.04 | **0.92** | −0.17 | −0.12 |
| *Strong* | −0.08 | 0.11 | **−0.97** | 0.09 | −0.11 | 0.18 | **−0.93** | 0.17 |
| *Feminine* | 0.17 | 0.16 | **0.72** | 0.63 | 0.12 | 0.17 | **0.78** | 0.53 |
| *Youthful* | 0.43 | −0.39 | −0.09 | **0.78** | 0.31 | −0.26 | −0.01 | **0.88** |

#### Supplemental Figure Legends

**Figure S1.** Reliability assessment of social trait ratings. **(A)** Number of online raters per trait. **(B, C)** Inter-rater consistency. Inter-rater consistency of each trait was estimated using **(B)** the intraclass correlation coefficient (ICC) (McGraw & Wong 1996) and **(C)** the Spearman's correlation coefficient ($\rho$). Inter-rater consistency was first calculated between raters and averaged within each module, and then averaged across modules. Error bars denote ±SEM across modules. **(B)** Intraclass correlation coefficient (ICC). **(C)** Spearman's ρ. **(D)** Correlation between ratings from different modules (i.e., different face images of the same identity). **(E)** Correlation between social traits. Color coding shows the Pearson's correlation coefficient *r*. **(F, G)** Correlation between patients' social trait ratings and the online social trait ratings by over 400 participants from the general population. **(F)** Individual patient (x-axis). Color coding shows the Spearman's correlation coefficient ρ. **(G)** Average across patients.

**Figure S2.** Dissimilarity matrices (DMs) calculated using face images. **(A)** The social trait DM. The DM was calculated using *z*-scored social trait ratings from the online participant group. **(B-D)** The neural response DMs. The DM was calculated using the normalized firing rate in all face-responsive neurons **(B)**, amygdala face-responsive neurons **(C)**, and hippocampal face-responsive neurons **(D)**. Color coding shows dissimilarity values. The social trait DM was correlated with the neural response DMs (permutation $P < 0.001$ for all face-responsive neurons, permutation $P = 0.008$ for all amygdala face-responsive neurons, and permutation $P < 0.001$ for all hippocampal face-responsive neurons).

**Figure S3.** Encoding and decoding models. **(A)** The percentage of neurons that significantly encoded a social trait. The dashed line indicates the chance level. **(B-D)** Average correlation coefficient across neurons for each social trait. Correlation was calculated between face identities. **(B)** All neurons. **(C)** All amygdala neurons. **(D)** All hippocampal neurons. **(E-H)** Average correlation coefficient across neurons for each social trait. Correlation was calculated between face images. **(E)** Face-responsive neurons. **(F)** All neurons. **(G)** All amygdala neurons. **(H)** All hippocampal neurons. Error bars denote ±SEM across neurons. Asterisks indicate a significant difference from 0 (two-tailed paired *t*-test). *: P < 0.05, **: P < 0.01, ***: P < 0.001, and ****: P < 0.0001. **(I, J)** Decoding social trait ratings across face identities using partial least squares (PLS) regression **(I)** and regression with principal component analysis (PCA) of neural responses **(J)**. Model predictability was assessed using the Pearson correlation between the predicted and actual trait ratings in the test dataset. The magenta bars show the observed response and the gray bars show the permuted response. Error bars denote ±SEM across permutation runs. Asterisks indicate a significant decoding performance (two-tailed two-sample *t*-test between observed vs. permuted). **: P < 0.01 and ****: P < 0.0001.

**Figure S4.** Temporal dynamics of neural encoding for each social trait. Blue: all amygdala neurons. Green: all hippocampal neurons. Error bars denote ±SEM across neurons. Asterisks shown on the top indicate a significant difference between amygdala and hippocampal neurons (two-tailed unpaired *t*-test, P < 0.05, corrected for multiple comparisons using false discovery rate [FDR] Q < 0.05).

#### Figure S1

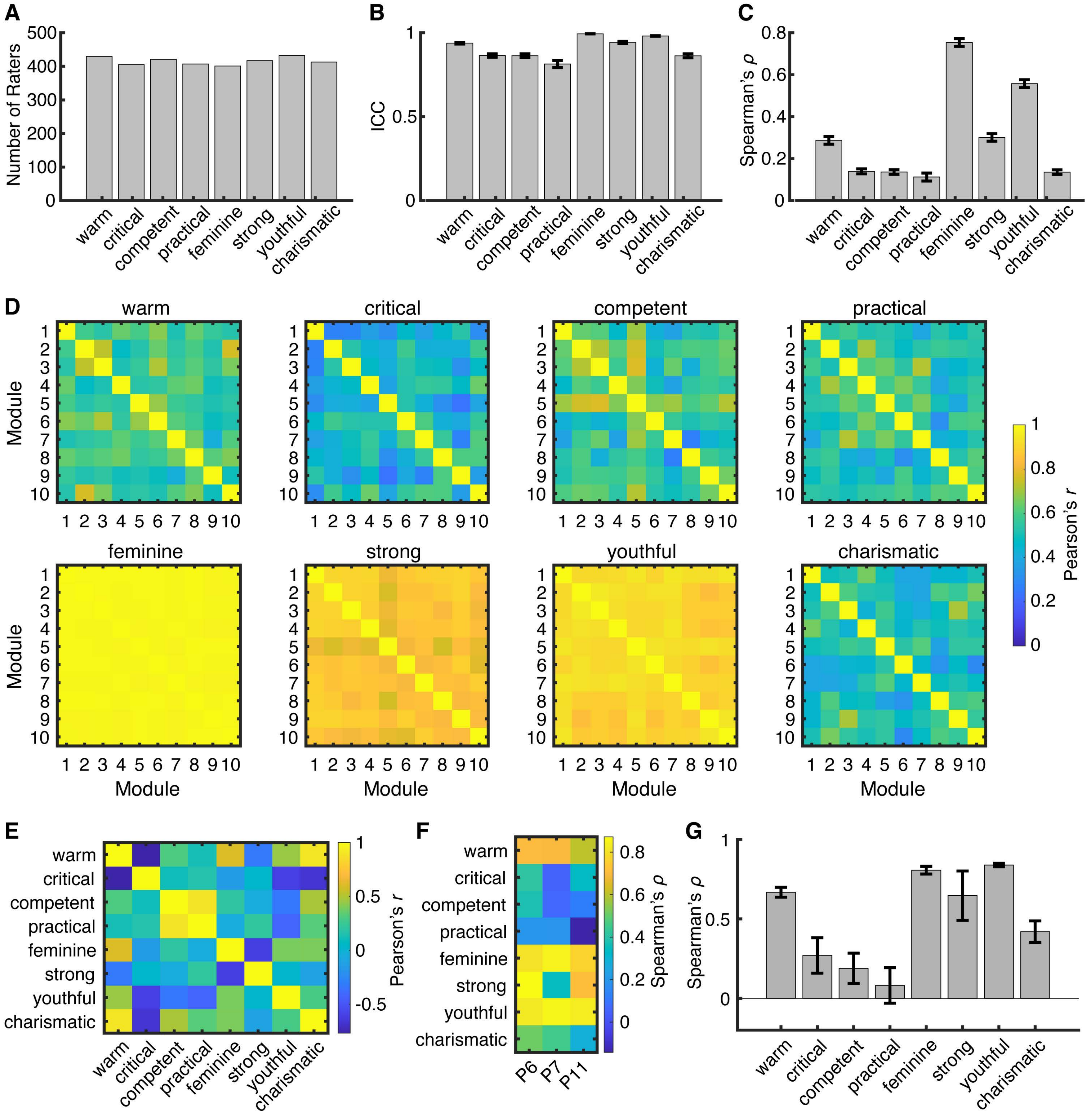
A
Number of Raters
warm
critical
competent
practical
feminine
strong
youthful
charismatic
B
ICC
C
Spearman's ρ
D
warm
critical
competent
practical
feminine
strong
youthful
charismatic
Module
Pearson's r
E
Pearson's r
F
Spearman's ρ
P6
P7
P11
G
Spearman's ρ

**Figure S2**

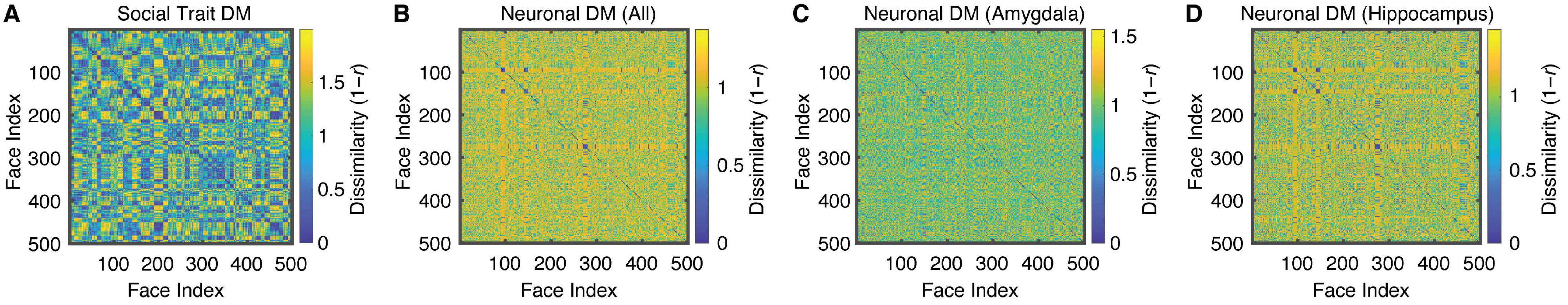

A
Social Trait DM
B
Neuronal DM (All)
C
Neuronal DM (Amygdala)
D
Neuronal DM (Hippocampus)
Face Index
Dissimilarity (1−r)
100
200
300
400
500
0
0.5
1
1.5

### Figure S3

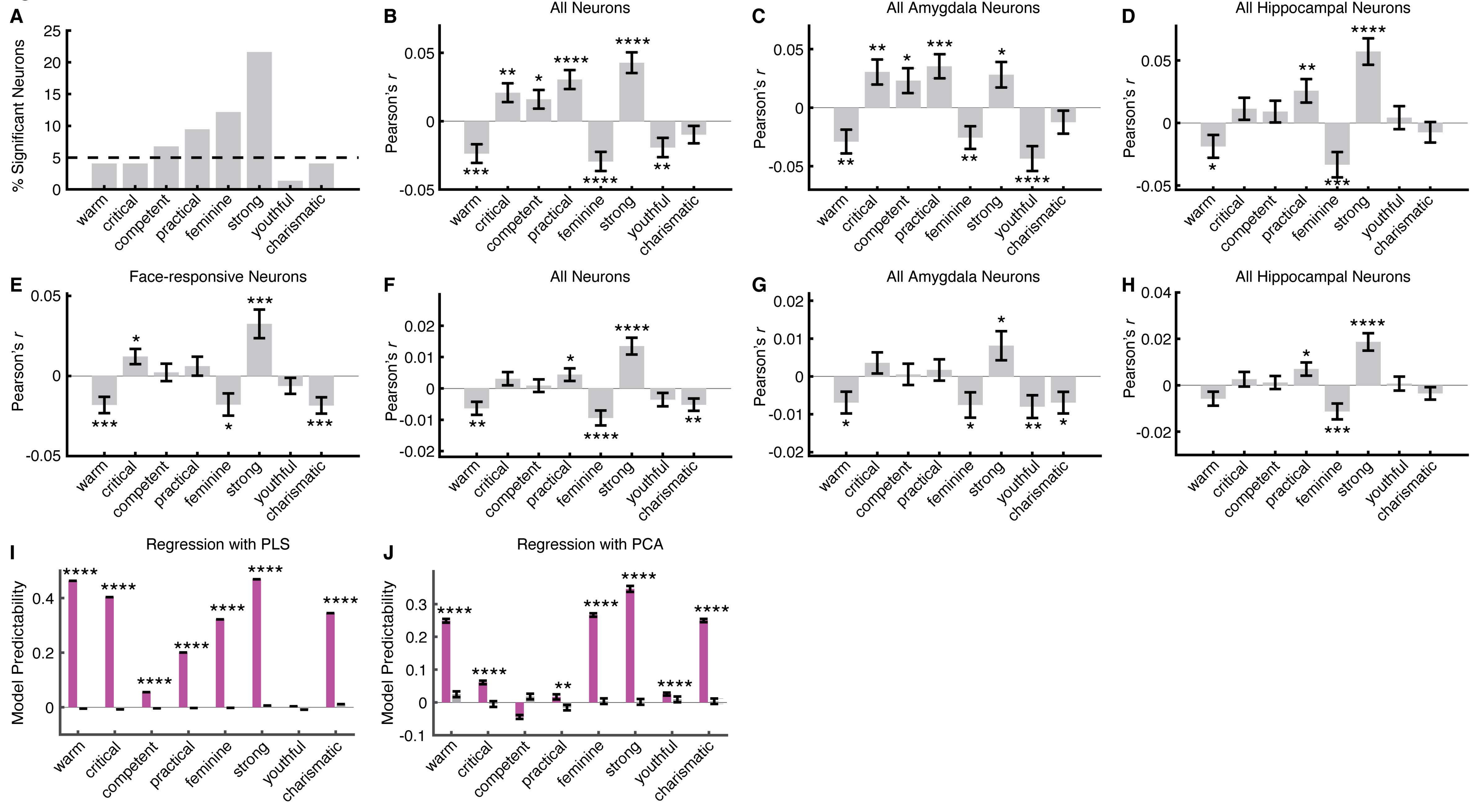

A
% Significant Neurons
warm critical competent practical feminine strong youthful charismatic
B All Neurons
C All Amygdala Neurons
D All Hippocampal Neurons
Pearson's r
E Face-responsive Neurons
F All Neurons
G All Amygdala Neurons
H All Hippocampal Neurons
I Regression with PLS
J Regression with PCA
Model Predictability

**Figure S4**

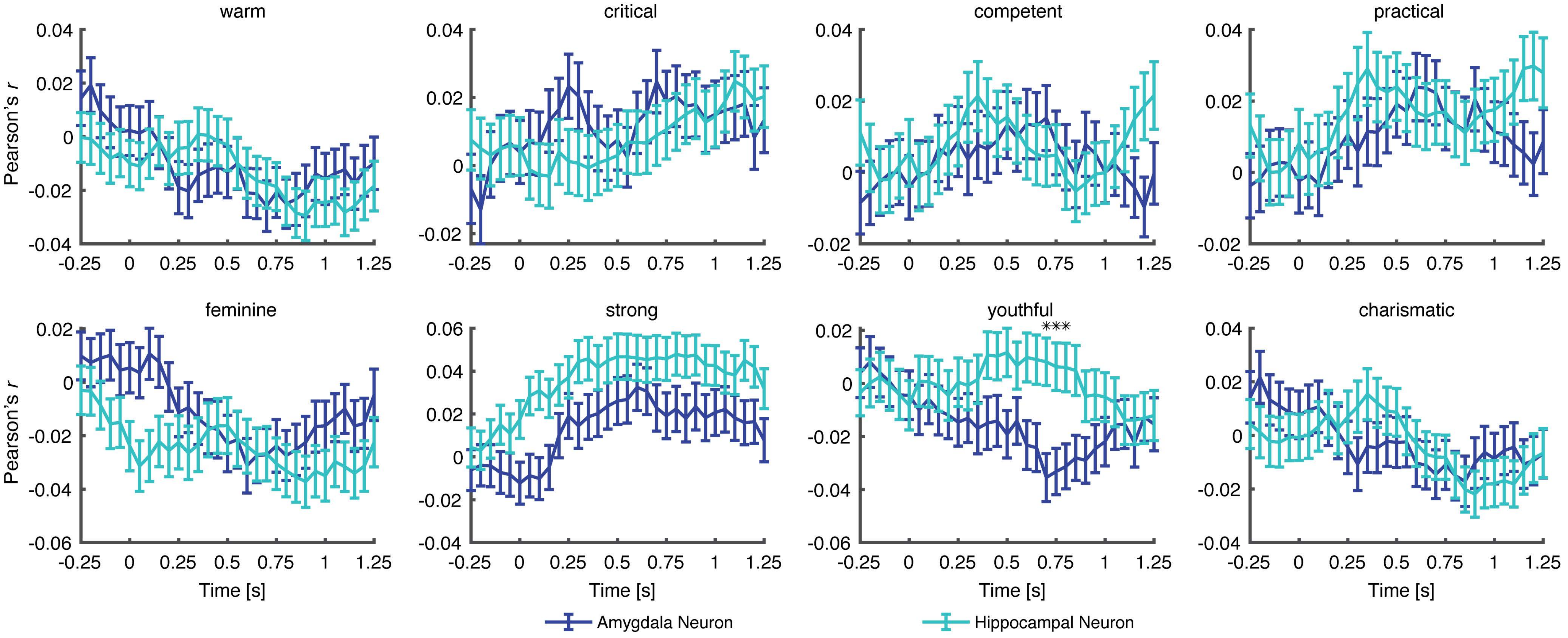
warm
critical
competent
practical
feminine
strong
youthful
charismatic
***
Pearson's r
Time [s]
Amygdala Neuron
Hippocampal Neuron
